# Structural Disconnection of the Posterior Medial Frontal Cortex Reduces Speech Error Monitoring

**DOI:** 10.1101/2021.09.13.460087

**Authors:** Joshua D. McCall, J. Vivian Dickens, Ayan S. Mandal, Andrew T. DeMarco, Mackenzie E. Fama, Elizabeth H. Lacey, Apoorva Kelkar, John D. Medaglia, Peter E. Turkeltaub

**Author notes:** **Corresponding author:** Peter E Turkeltaub MD, PhD, Address: Center for Brain Plasticity and Recovery and Neurology Department, Georgetown University Medical Center, Washington, District of Columbia, USA.

## Abstract

Optimal performance in any task relies on the ability to detect and repair errors. The anterior cingulate cortex and the broader posterior medial frontal cortex (pMFC) are active during error processing. However, it is unclear whether damage to the pMFC impairs error monitoring. We hypothesized that successful error monitoring critically relies on connections between the pMFC and broader cortical networks involved in executive functions and the task being monitored. We tested this hypothesis in the context of speech error monitoring in people with post-stroke aphasia. Diffusion weighted images were collected in 51 adults with chronic left-hemisphere stroke and 37 age-matched control participants. Whole-brain connectomes were derived using constrained spherical deconvolution and anatomically-constrained probabilistic tractography. Support vector regressions identified white matter connections in which lost integrity in stroke survivors related to reduced error detection during confrontation naming. Lesioned connections to the bilateral pMFC were related to reduce error monitoring, including many connections to regions associated with speech production and executive function. We conclude that connections to the pMFC support error monitoring. Error monitoring in speech production is supported by the structural connectivity between the pMFC and regions involved in speech production and executive function. Interactions between pMFC and other taskrelevant processors may similarly be critical for error monitoring in other task contexts.

## 1. INTRODUCTION

The ability to detect our own errors (i.e., error monitoring) is essential in daily life. The most well-documented neural correlate for error monitoring is the anterior cingulate cortex (ACC) (Botvinick et al., 1999; C. S. Carter et al., 1998; Gauvin et al., 2016). Notably, the neural activity associated with errors often extends to ACC-adjacent regions including the posterior superior frontal gyrus (pSFG) and medial precentral gyrus (mPG), which combined with the ACC, encompass a territory called the posterior medial frontal cortex (pMFC) (Ridderinkhof et al., 2007). Functional neuroimaging studies (i.e. using fMRI, EEG, PET) consistently demonstrate pMFC activity when individuals make errors on nonverbal tasks such as the Flanker task (Ullsperger & von Cramon, 2004) and verbal tasks such as tongue twisters (Gauvin et al., 2016). Various proposals have suggested that the pMFC, or parts of it, compute conflict, evaluate predicted outcomes, and compute the expected need to recruit control (Botvinick et al., 2001; Brown, 2013; Shenhav et al., 2013). However, our understanding of the pMFC’s role is limited because functional neuroimaging evidence allows for epiphenomenal explanations (e.g., that the pMFC supports error-related processing but not error monitoring per-se). In order to refute such epiphenomenal hypotheses, complementary evidence is needed to demonstrate that perturbation of the pMFC impairs error monitoring. Lesion studies are the most extreme case of perturbation to the pMFC and thus could provide converging evidence for non-epiphenomenal interpretations to this brain region.

To that end, there have been several small lesion studies concerning the ACC, a subregion of the pMFC. These lesion studies include case series that use nonverbal tasks to probe executive functions as well as conflict processing and error monitoring (Løvstad et al., 2012). Both conflict processing and error monitoring are often considered executive functions (e.g., Best & Miller, 2010), and are considered executive functions here as well. In sum, the available evidence has not comprehensively supported relationships between damage to the ACC and impairments of any executive functions. While one study (n=8) found that individuals with ACC lesions were not able to monitor task conflict, another study (n=4) found that individuals with ACC lesions monitored task conflict normally, and a third study (n=2) found that a participant with a left ACC lesion did not monitor task conflict but that a participant with a right ACC lesion monitored task conflict normally (di Pellegrino et al., 2007; Fellows & Farah, 2005; Swick & Jovanovic, 2002). There are also inconsistent results as to the consequences of ACC lesions on error monitoring as measured by post-error slowing. While the participants in Di Pellegrino et. al., 2007 (n=8) performing a Simon task did not display post-error slowing, participants in in Fellows and Farah 2004 (n=4) performing Stroop and Go No Go Tasks did display post-error slowing.

Intriguingly, an electroencephalography study of 5 individuals with large lesions encompassing the pMFC found an absence of the error-related negativity but intact error monitoring behavior on a Flanker task (Stemmer et al., 2004). In terms of other executive functions, two studies (n=4; n=2) found that individuals with ACC lesions had normal performance across a battery of executive functions (Baird et al., 2006; Fellows & Farah, 2005).

These lesion studies have been limited by only considering direct damage to the pMFC, which is relatively uncommon in stroke (Arboix et al., 2009), the predominant human lesion model. If the pMFC serves as a domain general error monitoring system, then it must interact with other brain structures that process the task-relevant information to be monitored. Thus, we hypothesize that disconnections between the domain general pMFC and task-relevant brain regions due to lesions should reduce error monitoring. We refer to this below as the “Disconnected Monitoring Hypothesis”.

Speech production is an ideal task to investigate the Disconnected Monitoring Hypothesis. Brain regions involved in speech processing are reasonably well-specified (e.g., Hickok & Poeppel, 2007; Rauschecker & Scott, 2009). Aphasia, an impairment of communication caused by brain damage, is common, and is often accompanied by a reduction in speech error monitoring (Robert C. Marshall & Tompkins, 1982; Oomen et al., 2001). Furthermore, a contemporary model of speech error monitoring allows for testable predictions about the role of communication between domain-general pMFC and task-relevant brain regions (Nozari et al., 2011). In their Conflict-Based Account, Nozari et. al. (2011) posits that the ACC (a subregion of the pMFC) monitors conflict that arises during activation of semantic and phonological representations during speech production. Correspondingly, our Disconnected Monitoring Hypothesis predicts that reduced speech error monitoring in aphasia results from structural disconnections between domain-general pMFC brain regions and brain regions that support speech production. Investigating critical brain structures and connections for speech error monitoring in aphasia also has clinical importance, and can potentially inform prognosis and future therapeutic approaches.

Only two studies to date have systematically investigated the neural correlates of speech-error monitoring in aphasia. One study found an electroencephalographic signature during incorrect naming trials centered over the pMFC (Riès et al., 2013), suggesting that the pMFC performs error-related processing in aphasia. In the second study, voxel-based lesion-symptom mapping found that frontal white matter lesions relate to reduced speech-error monitoring (Mandal et al., 2020). The association of lesioned white matter with reduced speech-error monitoring suggests that disconnections between certain brain regions may be related to poor error monitoring. However, this prior study did not examine disconnections directly, which would be critical to determine if disconnection of the pMFC from speech processing regions reduces speech error monitoring, as predicted by the Disconnected Monitoring Hypothesis.

Here, we evaluated the role of structural disconnections to the pMFC in error-monitoring in aphasia. We analyzed data from a subgroup of participants in a previous lesion-symptom mapping study (Mandal et. al. 2020) for whom diffusion-weighted images were available. We used support vector regression connectome-based lesion symptom mapping to identify white matter connections in which lost integrity of the connection, compared to controls, is related to reduced error monitoring. We hypothesized that disconnections between the pMFC and regions involved in speech processing would relate to reduced error monitoring.

## 2. MATERIALS AND METHODS

### 2.1 Participants

Data in this study was compiled from two studies including a clinical trial on transcranial direct current stimulation for aphasia (Cohort 1), and an investigation on inner speech in aphasia (Cohort 2). Many participants were part of both cohorts (n=22). The error-monitoring data from the combined cohorts is a subset from those used in the analyses in Mandal et al. 2020. Participants with left hemisphere stroke (n=51) in the present study were native English speakers, participated at least 6 months after their stroke, had no additional neurological or neuropsychiatric disorders, and produced enough errors to be able to measure error detection (see Behavioral Methods below). Controls with no history of stroke were matched to the stroke group on age and education (n=37). Controls were included to establish baseline connectome values to compare to the stroke group. All participants provided written informed consent. This research was approved by the Georgetown University Institutional Review Board.

### 2.2 Behavioral Methods

Speech errors and error detections were coded in the participants with left hemisphere stroke, and not the matched controls. The coding of speech errors and error detection in this participant pool has been previously detailed (Mandal et al., 2020) and will be described again below.

#### 2.2.1 Picture Naming Tasks

Participants with left hemisphere stroke completed picture naming tasks in which they name aloud black and white drawings presented one at a time. Participants in Cohort 1 received a 60-item version of the Philadelphia Naming Test (PNT) (Roach et al 1996). Participants in Cohort 2 received a total of 120 items across two 60-item naming tasks across two separate sessions: the 60-item version of the PNT, plus an additional 60 items which were normed in-house (stimuli available at https://www.cognitiverecoverylab.com/researchers). The mean interval between administration of 60 item naming tasks in Cohort 2 was 11 days. In sum, participants in Cohort 1 received 60 items, participants in Cohort 2 received 120 items, and participants in both Cohort 1 and Cohort 2 received a total of 180 items. In regards to the participants who were in Cohort 1 and 2, responses on all 180 items were included in this study. All picture naming responses underwent two types of coding: first, error coding, and second, error detection coding.

#### 2.2.2 Error Coding for Picture Naming

All spoken naming responses were recorded on video for offline coding of error types and error detection. Error coding rules paralleled those for the PNT (Roach et al., 1996). Errors were coded as phonological when the naming attempt shared either at least two phonemes, the stressed vowel, or first or last phonemes with the target. Errors were coded as semantic when the naming attempt was semantically related to the target. When the naming attempt was both phonologically and semantically related to the target, then the error was coded as “mixed” and was not considered as phonological or semantic.

#### 2.2.3 Error Detection Coding for Picture Naming

Detection coding was adapted from the protocol used by Schwartz and colleagues (e.g., Schwartz et al., 2016). Only the first naming attempt was considered for detection scoring. Error detection was tabulated separately for each error committed. Trials in which the participant made no response were not considered for detection scoring. Detections were coded when participants verbally indicated awareness of error commission (e.g., “dog….no that’s not right!”) or attempted to self-correct their error (e.g., “dog….cat!”). While participants had the opportunity to detect and correct their errors, they were not explicitly directed to judge the accuracy of their response on each trial. Since individual error detection rates cannot be measured when very few errors are committed, the detection rate for each error type was only analyzed in participants who committed at least 5 of the respective error type. Ultimately, 51 individuals committed at least 5 total errors, 41 individuals committed at least 5 phonological errors and 25 individuals committed at least 5 semantic errors. Semantic error detection rate was not considered for the present study because sample sizes of 30 and below can be underpowered for lesion-symptom mapping (Lorca-Puls et al., 2018). Total error detection rate was calculated as the total count of detected errors divided by the total count of all errors committed, excluding trials for which no response was given. Phonological error detection rate was calculated as the count of detected phonological errors divided by the count of phonological errors (Table 1).

**Table 1.** Participant Demographics

|  | Stroke Survivors:<br>Total Error Detection<br>Group | Stroke Survivors:<br>Phonological Error<br>Detection<br>Group | Matched Controls |
| --- | --- | --- | --- |
| Sample Size | 51 | 41 | 37 |
| Age (years) | 60.5 (9.5) | 60.5 (9.0) | 57.6 (12.6) |
| Sex (M/F) | 36 M,15 F | 29 M,12 F | 20 M,17 F |
| Education (years) | 16.2 (3.17) | 16.2 (3.16) | 16.9 (2.53) |
| Time Since Stroke (months) | 53.2 [6.2-256] | 49.2 [6.2-256] | NA |
| Lesion Size (cm<sup>3</sup>) | 110.1 (84.94) | 117.3 (80.89) | NA |
| WAB Yes/No<br>(Percent of Maximum Score) | 93% (7.04%) | 92% (7.19%) | NA |
| Mean Length of Utterance | 4.1 (2.2) | 3.9 (2.1) | NA |
| Naming Accuracy | 0.58 (0.3) | 0.52 (0.28) | NA |
| Naming No Responses ÷ Total Items | 0.098 (0.17) | 0.100 (0.16) | NA |
| Phonological Errors ÷ Total Errors | 0.484 (0.204) | 0.513 (0.201) | NA |
| Semantic Errors ÷ Total Errors | 0.166 (0.125) | 0.151 (0.109) | NA |
| Other Errors ÷ Total Errors | 0.350 (0.161) | 0.336 (0.161) | NA |
| Total Error Detection<br>(Percent of errors detected) | 41.3% (24.7%) | 41.6% (23.6%) | NA |
| Phonological Error Detection<br>(Percent of errors detected) | 39.6% (26.3%) | 39.6% (26.3%) | NA |
| Semantic Error Detection<br>(Percent of errors detected) | 36.1% (23.3%) | 36.9% (23.4%) | NA |

#### 2.2.4 Other Measures

Additional description of the participants is provided in the form of performance on tasks that demonstrate auditory comprehension and speech fluency (Table 1).

Auditory Comprehension: Participants completed the Auditory Verbal Comprehension subtest of the Western Aphasia Battery-Revised (WAB). Scores are reported from the Yes/No Questions task, where participants gave yes or no responses to 20 questions, which were either personal, environmental, or general (i.e., grammatically complex with no context) (Kertesz, 2007).

Speech Fluency: Mean Length of Utterance (MLU). Participants completed picture description tasks. The Mean Length of Utterance (MLU) was calculated from their description as the mean number of words used per utterance. Participants in Cohort 1 described the picture of a picnic scene in the WAB, whereas participants in Cohort 2 described the picture of a “Cookie Theft” in the Boston Diagnostic Aphasia Examination (Goodglass et al., 2000; Roth, 2011).

### 2.3 Imaging Methods and Connectome Construction

#### 2.3.1 Image acquisition

All brain images were acquired via a 3T Siemens Trio scanner at Georgetown via a 12-channel head coil. DWIs were acquired using single shot echo-planar imaging, consisting of 55 axial slices with a slice thickness of 2.5 mm, and voxel size of 2.5mm by 2.5 mm by 2.5 mm (repetition time (TR)= 7.5 s; echo time(TE) = 87 ms; field of view (FOV) = 240 mm × 240 mm; matrix size = 96 × 96; flip angle = 90°) (Figure 1.). In total, 80 volumes were acquired (60 at b = 1100 s/mm2, 10 at b = 300 s/mm2, 10 at b = 0 s/mm2).

**Figure 1.**
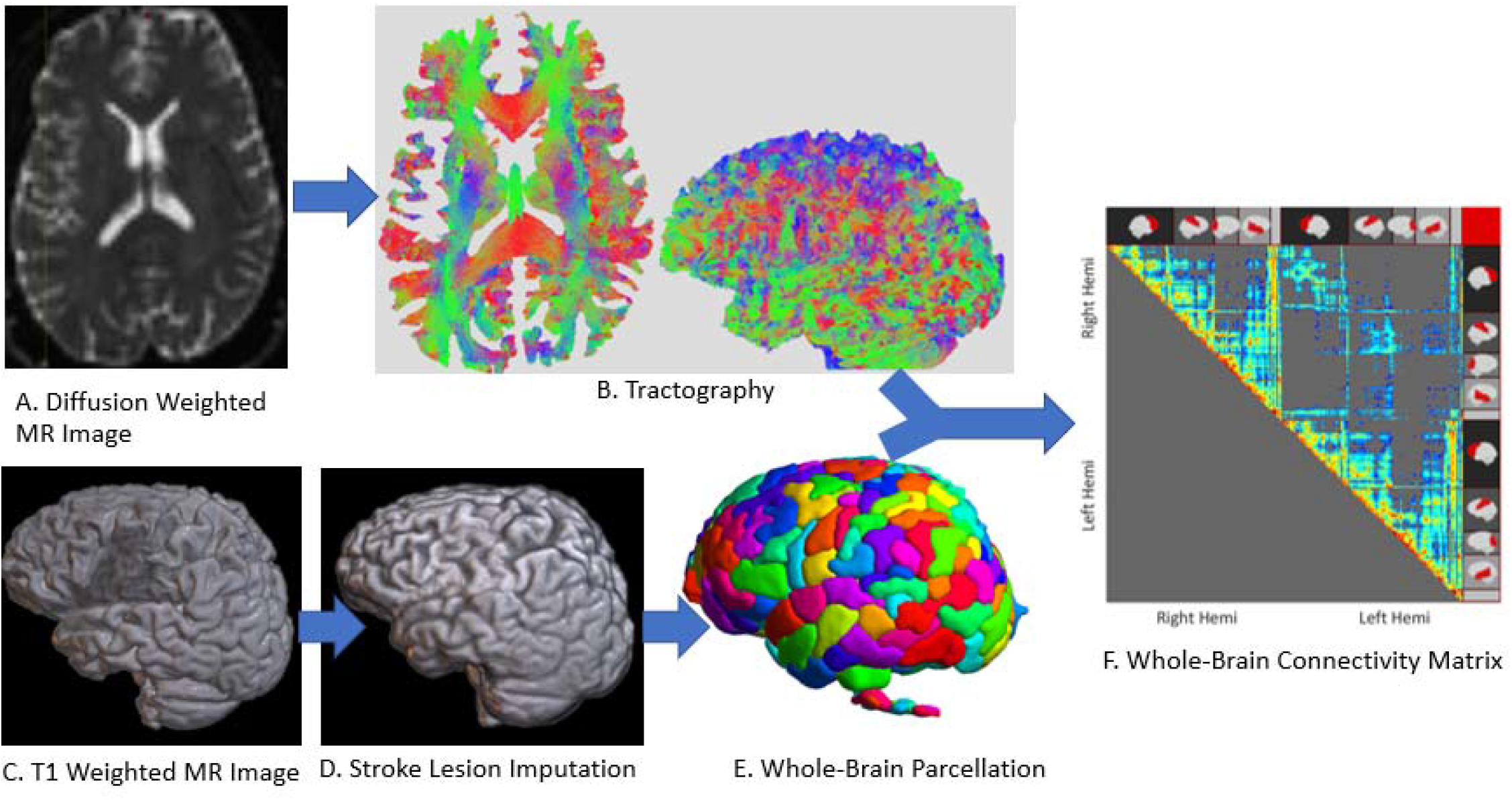
Pipeline of steps starting with diffusion weighted images (A), that are used to trace fiber tracts across the whole brain (B). Tracts are organized by which parcels of the brain they connect (E), ultimately resulting in a matrix diplaying connectivity between each parcel across the whole brain (F). Gray matter interface for the parcellation is determined based on T1-Weighted MR images (C), with the aid of imputation of tissue damaged by stroke (D).

Single volumes of Magnetization Prepared Rapid Acquisition Gradient Echo (MPRAGE) images were acquired and consisted of 176 sagittal slices with a slice thickness of 1 mm, and voxel size of 1mm by 1mm by 1mm (TR = 1900 ms; TE = 2.52 ms; inversion time= 900 ms FOV= 250 × 250 mm; matrix size= 256 × 256; flip angle = 9°).

#### 2.3.2 Construction of Structural Connectomes

In preparation for structural connectome construction, each stroke subject’s MPRAGE underwent an imputation process in order to enable tissue segmentation and brain parcellation. Imputation steps included automatic generation of the lesion in native space (Pustina et al., 2016), bias field correction, and skull-stripping of the brain. Lesioned voxels were filled by values from homotopic tissue values in the spared hemisphere. Additional lesion repair benefited from the fusion of T1 images from a group of at least 22 healthy matched controls warped to the lesioned brain via Advanced Normalization Tools (ANTs) (antsJointFusion) (Avants et al., 2011). Stroke and control subject MPRAGEs (imputed in stroke subjects) were submitted to FreeSurfer’s recon-all for cortical reconstruction (https://surfer.nmr.mgh.harvard.edu/). Lausanne atlas parcellations at scale 125 (Daducci et al., 2012) were generated from the output of recon-all with the preprocessed mean b0 image (see below) as the target image.

We derived structural connectomes from diffusion weighted images (DWIs) for stroke and control subjects using MRtrix 3.0 (Tournier et al., 2019) (Figure 1). Preprocessing steps for the DWIs included Gaussian noise removal (dwidenoise -extent 9,9,9), motion and eddy current correction (dwipreproc), and bias field correction (dwibiascorrect -ants). Voxel wise fiber orientation distributions were computed using multi-shell multi-tissue constrained spherical deconvolution (dwi2fod msmt_csd) based on a response function computed from the subject’s DWI data (dwi2response dhollander). (Jeurissen et al., 2014). For each individual, 15 million streamlines were generated by probabilistic anatomically-constrained tractography (Smith et al. 2012) on white matter fiber orientation distributions (tckgen -act, algorithm = iFOD2, step = 1, min/max length = 10/300, angle = 45, backtracking allowed, dynamic seeding, streamline endpoints cropped at grey matter-white matter interface). The five-tissue-type segmented image of the skull-stripped MPRAGE (imputed in stroke subjects) was warped into DWI space via ANTS and served as the anatomical image for anatomically-constrained probabilistic tractography. Spherical deconvolution informed filtering of tractograms 2 (SIFT-2) (R. E. Smith et al., 2015) was conducted in order to adjust streamline densities to be proportional to the underlying white matter fiber densities. Individual connections in the structural connectome were generated by assigning streamlines to parcels of the Lausanne atlas scale of 125. Network neuroscience research often refers to individual connections in the connectome as edges (Bassett & Sporns, 2017). However, since we are taking a theoretical approach that examines individual disconnections, we instead use the term “connections” throughout the paper.

Each streamline was multiplied by its respective cross-sectional multiplier derived via SIFT2, resulting in a value of apparent fiber density (AFD), which quantifies the relative cross-sectional area of the white matter fibers connecting two brain regions. The AFD may thus be thought of as the bandwidth of structural connectivity (i.e. fiber density) available between two brain regions. To enable group analyses, inter-subject AFD and connection density normalization was conducted (R. Smith et al., 2020). Specifically, each subject’s connectome was multiplied by the geometric mean of the ratio of the individual’s response function size at each b value to the group average response function size at each b value. Individual differences in white matter b0 intensity were accounted for by multiplying each connectome by the ratio of the mean median b0 value within the subject’s white matter mask to the grand mean median b0 value for the whole group. Inter-subject connection density normalization was then achieved through scalar multiplication of each connectome by the subject’s “proportionality coefficient” derived by SIFT2, denoted by μ, which represents the estimated fiber volume per unit length contributed by each streamline. Overall, each connection in the connectome quantifies the AFD of white matter tracts connecting two brain regions.

### 2.4 Experimental Design and Statistical Analysis

#### 2.4.1 Connectome-based Lesion-Symptom Mapping Analysis

Support vector regressions (SVR) were run to identify lesioned connections that cause reduced error detection rates. A connection was considered lesioned in a stroke participant when the AFD value was less than all of the control participants’ values at that connection. This binary definition of lesion includes connections with smaller values in stroke participants than controls, as well as those that are absent in stroke participants but detected in controls. This lesion definition permits a simple interpretation as it focuses on connections that were clearly lesioned by stroke.

The analyses were run using a method parallel to support vector regression-based lesion-symptom mapping (SVR-LSM) (e.g., DeMarco & Turkeltaub, 2018), where features based on brain imaging are used to predict a behavioral score. The key difference between connectome-based lesion-symptom mapping (e.g., Gleichgerrcht et al., 2017) and SVR-LSM is that brain-based features in SVR-LSM involve the lesion status of each voxel across structural images, whereas the brain-based features in this connectome symptom mapping analysis are the lesion status of each structural connection across connectomes. Since the presence or absence of individual small connections may vary across individuals irrespective of lesions, only connections present in 100% of control subject connectomes (n=37) were included in the analyses. To avoid spurious relationships based on connections that are only lesioned in one or a few subjects or happen to be smaller than controls based on chance alone, the analysis only considered connections that were lesioned in at least 20% of participants with left-hemisphere stroke.

The dependent variable modeled by SVR is percent detection rate on errors made during picture naming. Lesion volume in voxels was covaried out of both connectome values and detection rate scores prior to modeling. SVR hyperparameters included Cost (set to MATLAB’s default), kernel (radial basis function), and kernel scale (set to 1).

Since lesion-mapping efforts have found that detection of phonological errors has stronger lesion-deficit associations than detection across all errors (Mandal et al., 2020), two separate analyses were run for detection across all errors (n=51) and for only phonological errors (n=41). Resulting SVR beta weights for each connection were assigned p values using permutation tests, in which error detection rates were randomly assigned to connectomes 10,000 times, and each connection was ranked among the 10,000 permutation-based values for that connection. Statistical thresholding used a continuous familywise error rate (CFWER) method, with a family-wise error rate of .05 and v ranging from 1 to 20 (Mirman et al., 2018). In CFWER, the v^th^ most maximal test statistic from each permutation is recorded to form a null distribution, which is then used to provide an adjusted p-value threshold that limits the number of expected false-positive results to v. For example, at v=20 CFWER produces a p-value threshold at which there is a 5% chance of obtaining 20 significant connections in the entire connectome. Therefore, analyses that result in 20 or more significant connections can be interpreted as non-random. The confidence in individual connection-wise results increases with the number of connections that survive above the v value. Whereas full family-wise error correction provides greater confidence for interpreting the importance of individual connections, it may be overly strict for maps in which loss of individual connections do not produce a dramatic behavioral impairment because the behavior relies on a network involving multiple connections. Analyses were run with v values ranging from 1 to 20 to increase power to detect maps involving over 20 connections.

## 3. RESULTS

### 3.1 Behavioral Scores

#### 3.1.1 Total Error Detection Group

A total of 51 participants were included in the SVR analysis on detection rate across all errors. Phonological errors accounted for 48.4% of all errors in these participants. On average, detection rate across all errors was 41.3% (*SD* = 24.7%) (Table 1).

#### 3.1.2 Phonological Error Detection Group

A total of 41 participants were included in the SVR analysis on detection rate across phonological errors. On average, detection rate across phonological errors was 39.6% (SD = 26.3%) (Table 1).

### 3.2 Tractography

The tractography revealed expected patterns in the connectivity between brain parcels in control participants, including a greater presence of connections between adjacent cortical regions, subcortical-to-cortical connections, and interhemispheric connections that are roughly homotopic (Figure 2).

**Figure 2.**
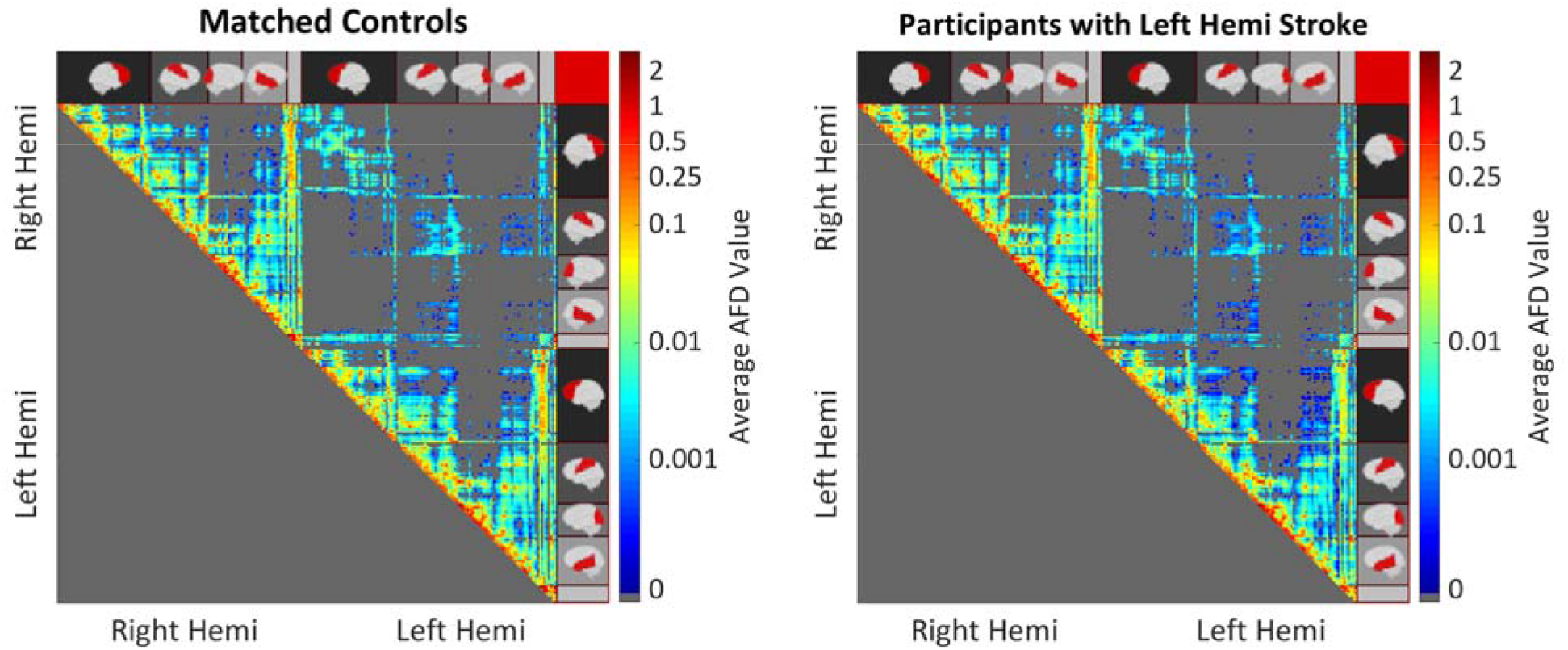
Mean AFD connection values for control participants (left) and participants with left hemisphere stroke (right). Left Hemi indicates Left Hemisphere, and Right Hemi indicates Right Hemisphere. Cortical regions are indicated on the sides of the matrix, with the empty gray bar indicating the subcortical regions (i.e. basal ganglia and brain stem). Connections between adjacent regions are represented on the main diagonal.

Lesion overlap maps (Figure 3c-3d) revealed broad coverage of disconnections within the left hemisphere, and between roughly homotopic regions of the left and right hemisphere. Connections that were lesioned in at least 20% of stroke participants were included in the analyses examining disconnections associated with reduced error detection (Figure 3e-3f).

**Figure 3.**
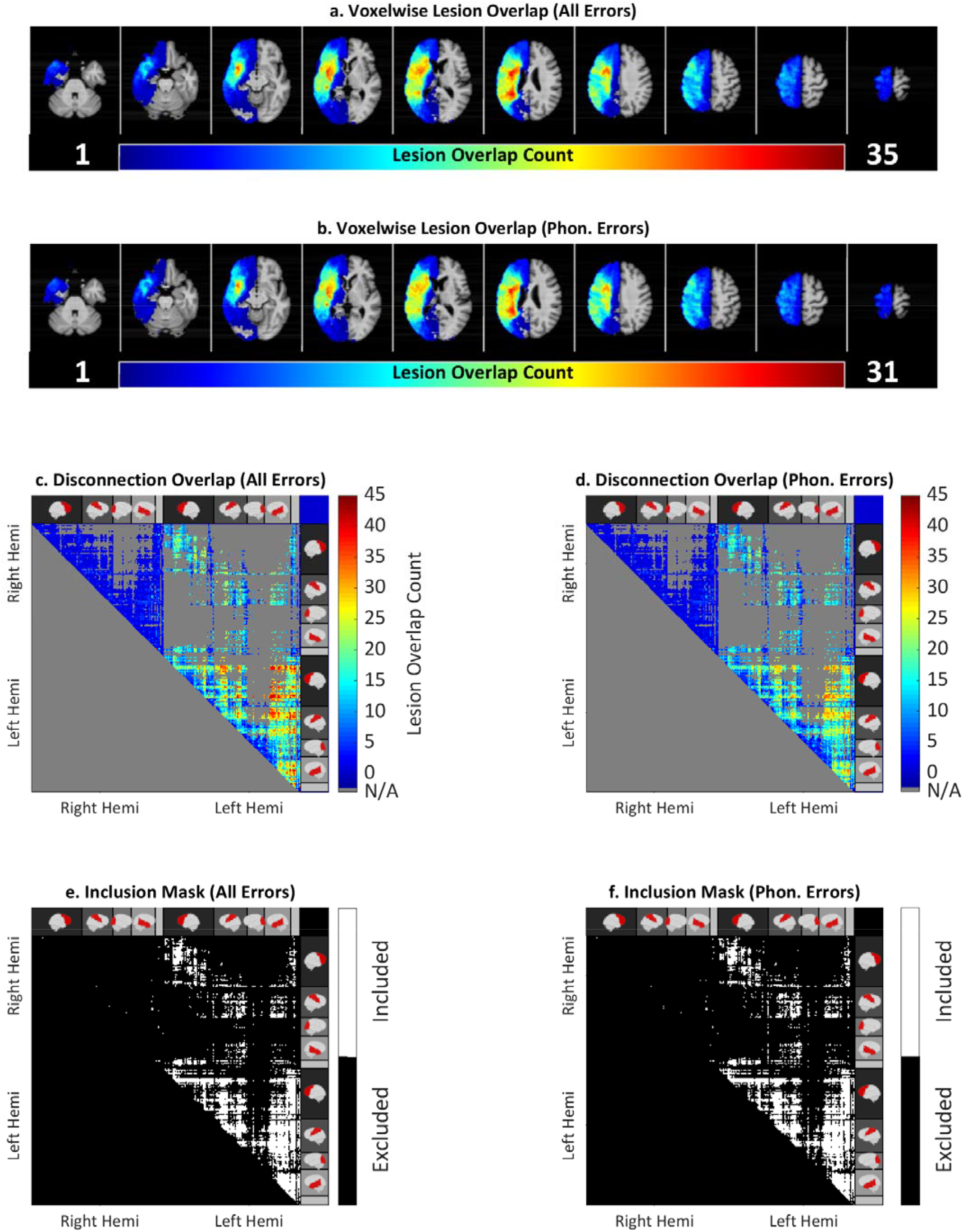
Top: Lesion overlap map identifying common locations of brain damage across each participant (a,b). Middle: Overlap maps counting total disconnections across participants included each analysis (c,d). Bottom: Maps of connections included for each analysis (e,f). All Errors indicates analysis for detection rate across all speech errors(N=51). Phon. Errors indicates analysis for detection rate across phonological speech errors(N=41). Right Hemi indicates Right Hemisphere, Left Hemi indicates Left Hemisphere.

### 3.3 Support Vector Regressions

#### 3.3.1 Detection Rate Across All Speech Errors

##### Interpretability of Significant Maps

In order to reveal connections that serve a general role in speech error monitoring, the first SVR analysis examined disconnections associated with reduced detection rate across all speech errors, irrespective of the type of error. Maps for detection rate across all errors were significant at thresholds v = 3, 5, 12, and 14. At v = 14, a total of 14 connections survived (Table 2, Figure 4). Although the maps are statistically significant at these thresholds, the interpretability of the individual connections are limited because the maps barely achieve the required number of connections at each threshold of v = 3, 5, 12, and 14. For example, there is a 5% chance that all 14 of the specific connections at v=14 occurred by chance. In consideration of this low level of confidence in the individual observed connections, only general trends across the map at v=14 are described below.

**Table 2.** Significant Connections for Detection Rate Across All Speech Errors. Values are reported in as averages, with standard deviations in parentheses, range in square brackets.

| Connection<br>(Parcel 1, Parcel 2) | MNI Coordinates<br>(Parcel 1, Parcel 2) | SVR *Beta*<br>Value | *p* Value | Significant<br>CWFER *v*<br>(min) |
| --- | --- | --- | --- | --- |
| Left parsopercularis 1, Left superiorfrontal 6 | (-38, 15, 12), (-19, 16, 51) | 8.62 | 0.0002 | 3 |
| Left parsopercularis 2, Left superiorfrontal 5 | (-47, 11, 11), (-7, 29, 54) | 8.84 | 0.0002 | 3 |
| Left parstriangularis 1, Left superiorfrontal 6 | (-40, 31, 2), (-19, 16, 51) | 9.03 | 0.0002 | 3 |
| Left caudalmiddlefrontal 1, Left insula 2 | (-35, 20, 47), (-37, -5, -2) | 8.16 | 0.0003 | 5 |
| Left parsopercularis 2, Left superiorfrontal 2 | (-47, 11, 11), (-9, 43, 17) | 7.84 | 0.0003 | 5 |
| Right caudalanteriorcingulate 1, Left insula 4 | (6, 17, 24), (-30, 12, 7) | 7.93 | 0.0003 | 5 |
| Left parsopercularis 1, Left superiorfrontal 8 | (-38, 15, 12), (-6, -1, 60) | 8.14 | 0.0005 | 12 |
| Left parsopercularis 2, Left superiorfrontal 6 | (-47, 11, 11), (-19, 16, 51) | 8.02 | 0.0005 | 12 |
| Left superiorfrontal 6, Left superiortemporal 5 | (-19, 16, 51), (-46, 8, -21) | 8.27 | 0.0005 | 12 |
| Left parsopercularis 1, Left caudalmiddlefrontal 1 | (-38, 15, 12), (-35, 20, 47) | 7.77 | 0.0006 | 12 |
| Left parsopercularis 1, Left superiorfrontal 5 | (-38, 15, 12), (-7, 29, 54) | 7.79 | 0.0006 | 12 |
| Left parstriangularis 1, Left superiorfrontal 5 | (-40, 31, 2), (-7, 29, 54) | 7.81 | 0.0006 | 12 |
| Left rostralmiddlefrontal 3, Left insula 2 | (-40, 39, 14), (-37, -5, -2) | 8.01 | 0.0007 | 14 |
| Right thalamusproper, Left precentral 6 | (14, -17, 6), (-40, -13, 33) | 8.34 | 0.0007 | 14 |

**Figure 4.**
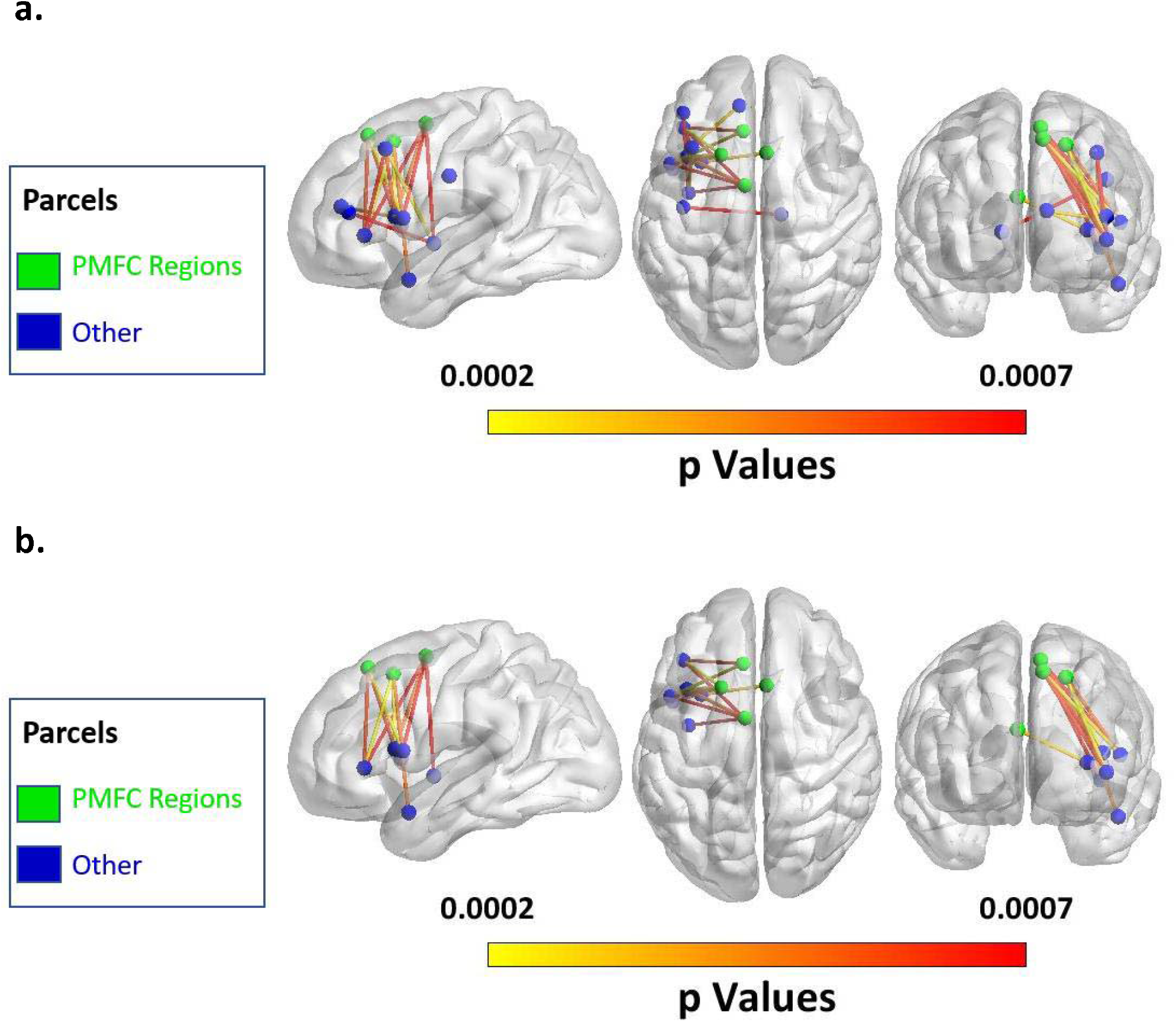
(a.) Map of all significant connections at v=14 for detection rate across all speech errors. (b.) Map of significant connections involving the posterior Medial Frontal Cortex (pMFC) at v=14 for detection rate across all speech errors

##### Significant Connections

Twelve of the 14 connections were within the left hemisphere, and the other two were interhemispheric. All 14 connections involved regions of the frontal lobe: with 9 connections within the frontal lobe, 3 connections between the frontal lobe and insula, 1 connection between the frontal and temporal lobe, and 1 connection between the frontal lobe and thalamus. As hypothesized, the majority of significant connections involved regions of the pMFC, including the ACC, and pSFG. (9 of 14; 63.4%; Figure 4b).

#### 3.3.2 Detection Rate Across Phonological Speech Errors

##### Interpretability of Significant Maps

In order to identify connections that are specifically involved in monitoring phonological aspects of speech, the next SVR analysis examined disconnections associated with reduced detection of phonological errors. Maps for detection rate across phonological speech errors were significantly non-random at all thresholds from v = 1 through v = 20. At v = 20, 46 connections survived thresholding (Table 3, Figure 5). Since the map surpasses the required number of connections at v=20, the interpretability for individual connections is strong. There is a 5% chance that 20 of the 46 significant connections occurred by chance. Therefore, we can consider individual significant connections at v=20 with confidence that they are nonrandom.

**Table 3.**
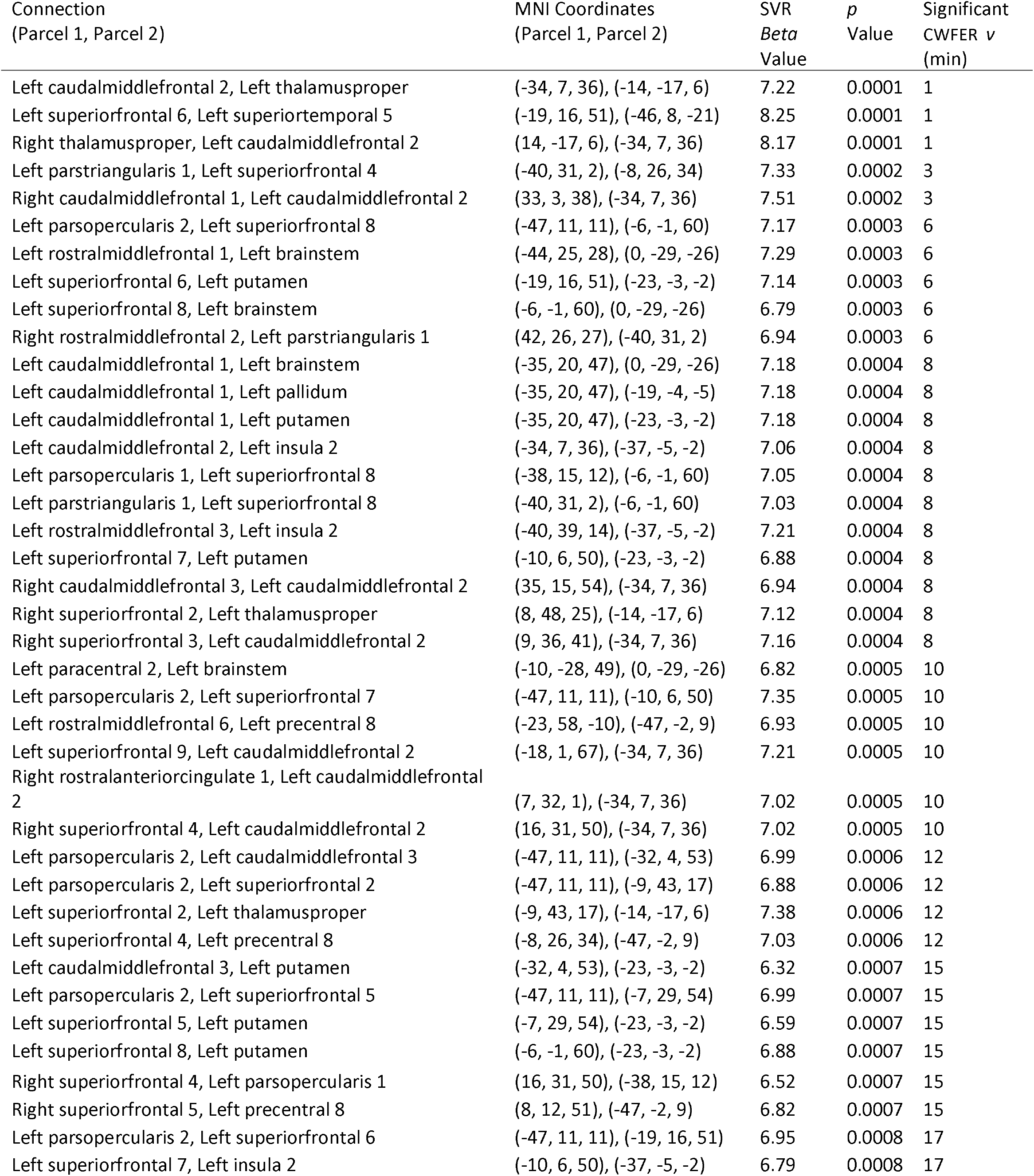
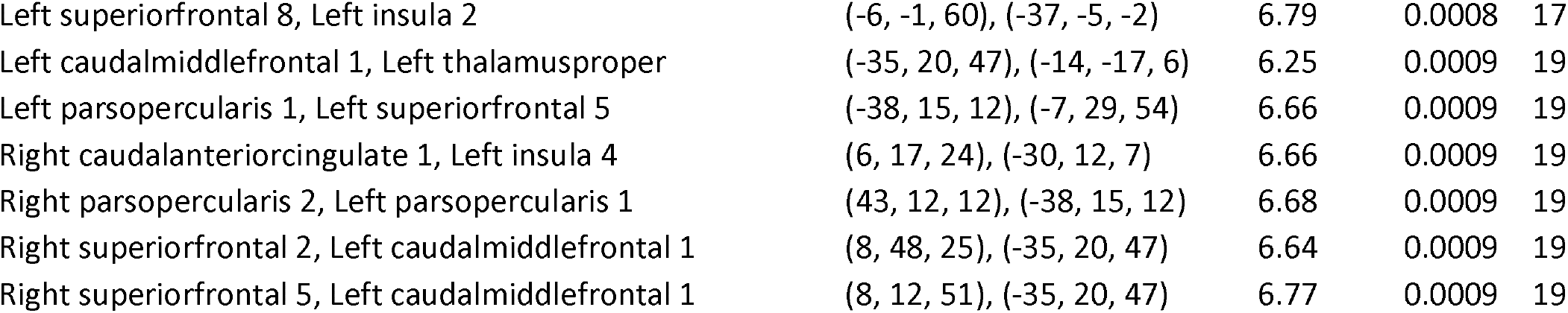
Significant Connections for Detection Rate Across Phonological Speech Errors.

**Figure 5.**
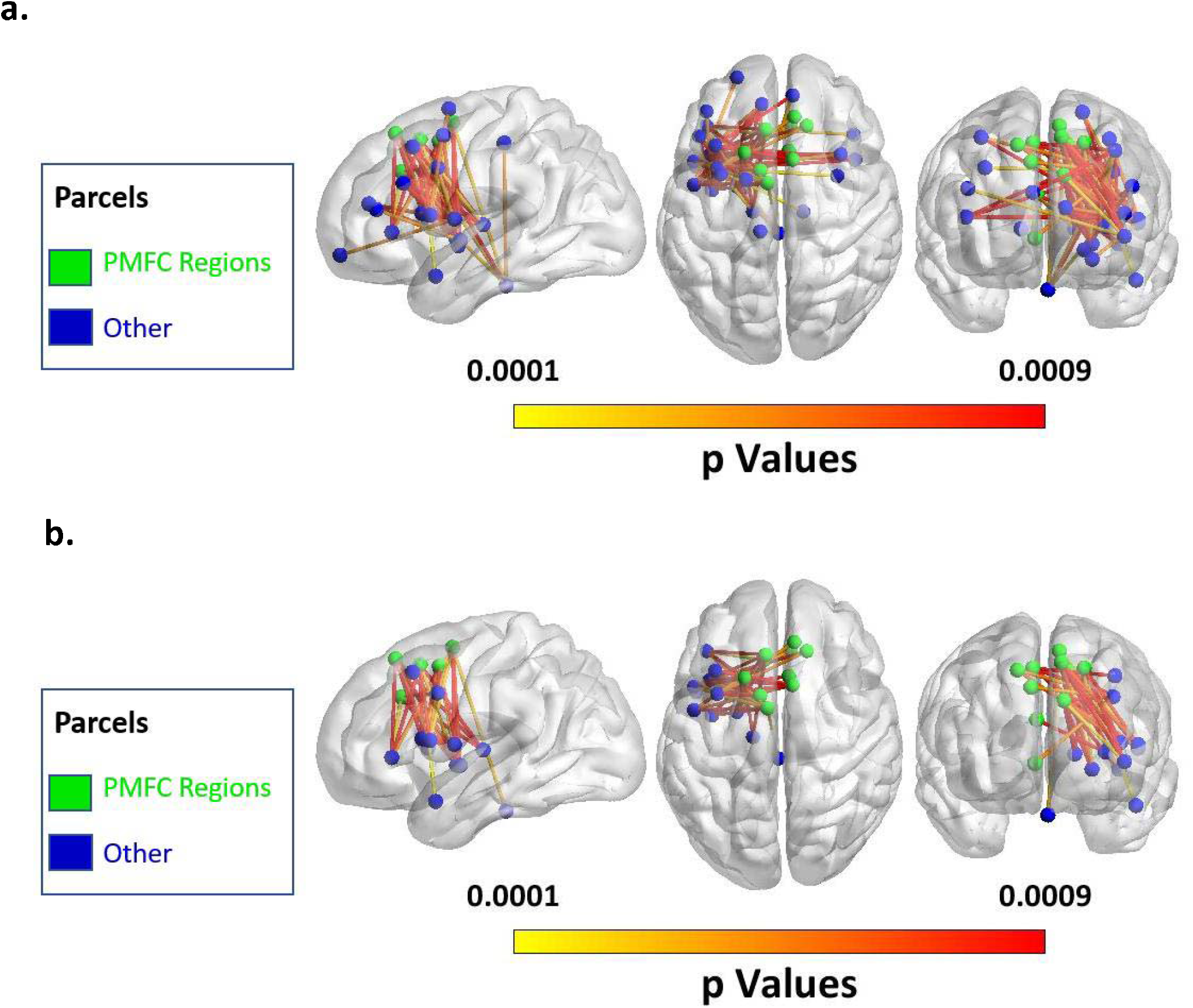
(a) Map of all significant connections at v=20 for detection rate across phonological speech errors. (b) Map of significant connections involving the pMFC at v=20 for detection rate across phonological speech errors

##### Significant Connections

Twenty-eight of the 46 connections were within the left hemisphere, and the other 14 were interhemispheric. Forty five of the 46 connections involved regions of the frontal lobe: with 24 connections within the frontal lobe, 5 connections between the frontal lobe and insula, 1 connection between the frontal and temporal lobe, and 15 connections between the frontal lobe and subcortical structures (i.e., thalamus, brainstem, basal ganglia). The only connection that did not involve the frontal lobe connected an area of the parietal lobe to the insula. As hypothesized, the majority of the connections involved regions of the pMFC including the ACC, and pSFG (24 of 46; 52.2%).

All 24 connections involving the pMFC connected the pMFC to regions of the brain’s left hemisphere (Figure 5b). The region with the most connections from the pMFC implicated in the analysis was the inferior frontal gyrus, with 9 significant connections (8 to pars opercularis, 1 to pars triangularis). Six significant pMFC connections were to subcortical regions (5 to putamen, 1 to brainstem). Five significant pMFC connections were to dorsolateral prefrontal cortex (dlPFC), and two each were to the ventral precentral gyrus and the insular cortex. One pMFC connection was to the anterior superior temporal gyrus.

## 4. DISCUSSION

### 4.1 Main findings

Our whole-brain CLSM analyses confirmed the Disconnected Monitoring Hypothesis, demonstrating that structural disconnections involving the pMFC are associated with reduced speech-error monitoring in aphasia. In the maps for detection rate across both total errors and phonological errors, over half of the disconnections implicated in reduced monitoring involved the pMFC. A growing body of functional neuroimaging evidence suggests that error monitoring recruits activity of the ACC and adjacent regions in the pMFC territory including the pSFG and mPG (Ridderinkhof, 2004). Our findings add complementary lesion-based evidence for this relationship, supporting the existence of an error-monitoring network that relies on the pMFC.

Furthermore, we found an additional network of regions where connections to the pMFC support error monitoring in speech. Specifically, we found a predominance of significant connections between the pMFC and regions known to support speech production including motor regions and the left inferior frontal gyrus (IFG) (Mirman et al., 2015; Pillay et al., 2017; Wilson, 2017). There were also significant connections between the pMFC and the dlPFC, a region canonically involved in executive function. This is the first study to present direct lesion-based evidence that structural connectivity between the pMFC and regions involved in speech production and executive function are important for error monitoring in speech.

### 4.2 The Role of the pMFC and Executive Regions in Error Monitoring

Only a few lesion studies with small sample sizes have examined the role of the pMFC in error monitoring, yielding mixed results (Baird et al., 2006; di Pellegrino et al., 2007; Fellows & Farah, 2005; Løvstad et al., 2012; Stemmer et al., 2004). The present study (n=51) demonstrates that damage to connections to the pMFC is associated with reduced error monitoring in speech. Regions across the pMFC were implicated including the ACC as well as the pSFG. It is curious why prior lesion studies on the role of the pMFC in error monitoring have yielded mixed findings. One possible cause of these mixed findings is that many of these studies examined lesions to the ACC, a subregion of the pMFC, and often only unilateral lesions to the ACC. If the role of the unilateral ACC is partially redundant with the contralateral ACC, or partially redundant with other subregions of the pMFC such as the contralateral or ipsilateral pSFG, then lesion-deficit relationships with the ACC could be more subtle. In that case, studies could include some participants with lesions to part of the pMFC that caused subtle changes in their error monitoring behavior and not obvious impairments. Such subtle lesion-deficit associations would require large sample sizes to consistently detect. Thus, partial redundancy between contralateral or ipsilateral pMFC regions in combination with small sample sizes could cause lesion studies on the pMFC to yield mixed findings. It is also notable that prior lesion studies examined error monitoring of cognitive control tasks, which are thought to rely on the bilateral prefrontal cortex (Miller & Cohen, 2001). One could imagine that error monitoring of ‘bilatera? cognitive control tasks readily recruits bilateral pMFC regions. This bilateral pMFC recruitment may foster partial redundancy that obscures lesion-deficit relationships. On the other hand, the present study examined monitoring during speech production, a task that relies on the brain’s left hemisphere(Szaflarski et al., 2006). Therefore, the present study may have benefited from extra sensitivity to lesion-deficit relationships by examining monitoring during a ‘unilateral’ task. Our disconnection-based approach may have garnered additional sensitivity because we examined error monitoring during a left-hemisphere task in participants who have disconnections involving the left hemisphere. In other words, we would expect disconnections involving the left hemisphere to strongly impact error monitoring because the left hemisphere contains the task-relevant information being monitored.

The pMFC and dlPFC have been previously theorized to communicate during error monitoring. For example, conflict monitoring theory proposes that the pMFC generates a conflict-based error signal and then recruits the dlPFC to exert executive control (C. S. Carter & van Veen, 2007; Kerns et al., 2004). Alternatively, the hierarchical error representation model proposes that the pMFC generates a prediction-based error signal that is sent to the dlPFC to hold error prediction representations in working memory (Alexander & Brown, 2015). Under the hierarchical error representation model, the dlPFC also signals to the pMFC in order to refine prediction-based error signaling (Alexander & Brown, 2015). Although the exact information being communicated remains under debate, the present study confirms that structural connectivity between the pMFC and dlPFC is important in error monitoring.

### 4.3 Connections Between pMFC and Speech Production Regions Support Error Detection

A current debate in the field of error monitoring concerns the degree to which the circuitry for error monitoring is specific to the task being monitored. This debate is especially active in the literature for speech error monitoring (Nozari, 2020; Roelofs, 2019). Since functional neuroimaging studies consistently find error-related pMFC activation across task domains (Gauvin et al., 2016; Ullsperger & von Cramon, 2004), we posited that the pMFC plays a domain-general processing role. We predicted that the pMFC would need to communicate with specific brain regions responsible for task-relevant representations in order to support error monitoring. Indeed, we found that speech error monitoring relies on connections between the pMFC and brain structures that process speech, particularly regions canonically associated with speech production.

Our finding of the importance of connections between the pMFC and regions supporting speech production is consistent with Nozari et. al.’s Conflict Based Account which proposes that error-monitoring relies on the interplay between speech production regions and general error monitoring machinery. The IFG is thought to support speech production, and was prominently involved in significant disconnections with the pMFC in our results. It is worth noting that the IFG has also been proposed to support a wide range of general executive functions that aid in language processing (Fedorenko et al., 2012; Fedorenko & Kanwisher, 2011; Lambon Ralph et al., 2017; Nozari et al., 2016; Thompson et al., 2018). Thus, an alternative interpretation of our results is that the connections between the IFG and pMFC are part of the general executive function circuity that supports error monitoring across task domains. However, connections between the pMFC and other regions thought to support speech production, including the ventral motor cortex, the basal ganglia and the anterior insula, were also implicated by our results (Dronkers, 1996; although see Fedorenko et al., 2015 regarding the role of anterior insula in speech production; Jürgens, 2002), strongly supporting the role of connections between speech production machinery and pMFC in speech error monitoring.

Ultimately, the task-specificity of the connections found in our study remains unclear because we did not compare speech error monitoring to error monitoring in other domains (e.g., nonverbal tasks). Within the context of speech error monitoring, the Disconnected Monitoring Hypothesis would predict that connections between the pMFC and different cortical regions might support monitoring of different types of errors. Unfortunately, we did not have enough semantic errors to directly compare disconnections that affect monitoring of phonological and semantic errors. Thus, our results do not provide clear evidence for the specificity of neural connections for error monitoring in specific task contexts. However, our results may indirectly imply such specificity because the results were stronger when we examined one specific error type (i.e., phonological errors) than when we examined all errors together. If error-monitoring relies on connections between the pMFC and processors specific to the signals being monitored, one would expect an analysis that mixes multiple error-types to yield relatively weaker results since individual connections may only be important for monitoring certain error types and not others. Consistent with this prediction, the pattern of connections observed for all errors was similar to that observed for phonological errors, likely reflecting the predominance of phonological errors in our sample, but the results were less robust when all errors types were examined together. Further research is needed to dissect the task-specificity of neural substrates that support error-monitoring.

### 4.4 Limitations

This study utilized a naturalistic measurement of error monitoring in the sense that participants were not explicitly directed to monitor their errors. The scoring of error monitoring therefore relied on participants spontaneously monitoring their errors. This study’s measurement method is standard in the field of speech error monitoring in aphasia, and is often used in research on the relationship between error monitoring and clinical outcomes (Robert C. Marshall et al., 1982, 1994; Nozari et al., 2011; Schwartz et al., 2016). However, we cannot preclude the possibility that participants successfully monitored some errors but chose not to display spontaneous error monitoring behavior. The inclusion of explicit instructions to monitor all responses for accuracy could reduce this possibility, but may affect the strategies and processes participants use to monitor their errors (e.g., Grützmann et al., 2014).

### 4.5 Conclusions

This connectome-based lesion analysis demonstrated that damage to connections involving the pMFC reduces the monitoring of speech errors. Specifically, damage to structural connections between the pMFC, other structures involved executive functions, and speech production regions led to a reduction in error detection, particularly for phonological errors. These findings are consistent with the notion that error monitoring critically relies on structural connectivity between task-specific cortical regions and domain-general control regions, including the pMFC. These results align with production-based models of speech error monitoring (Gauvin & Hartsuiker, 2020; e.g., Nozari et al., 2011). In clinical settings, damage to fiber tracts that connect to the pMFC may predict reduced error monitoring.

## Conflicts of Interest

The authors declare no competing financial interests.

## Funding

This work was supported by the National Institute on Deafness and other Communication Disorders [NIDCD grants R03DC014310 and R01DC014960 to PET, F31DC014875 to MEF, F30DC019024 to JDM, F30DC018215 to JVD], the Doris Duke Charitable Foundation [Grant 2012062 to PET], the National Center for Advancing Translational Science [KL2TR000102 to PET], the National Institute of Health’s StrokeNet [Grant U10NS086513 to ATD], and the National Center for Medical Rehabilitation Research [K12HD093427 to ATD].

## Acknowledgements

We are grateful for Zainab Anbari, Maryam Ghaleh, Mary Henderson, Harshini Pyata and Katherine Spiegel, for their contributions to data collection, as well as our participants for their time and effort in the study.

## Notes

### Competing Interest Statement

The authors have declared no competing interest.

